# Medium depth influences O_2_ availability and metabolism in cultured RPE cells

**DOI:** 10.1101/2023.03.01.530623

**Authors:** Daniel T. Hass, Qitao Zhang, Gillian A. Autterson, Richard A. Bryan, James B. Hurley, Jason ML. Miller

## Abstract

**Purpose:** RPE oxidative metabolism is critical for normal retinal function and is often studied in cell culture systems. Here, we show that conventional culture media volumes dramatically impact O_2_ availability, limiting oxidative metabolism. We suggest optimal conditions to ensure cultured RPE is in a normoxic environment permissive to oxidative metabolism.

**Methods:** We altered the availability of O_2_ to human primary RPE cultures directly via a hypoxia chamber or indirectly via the amount of medium over cells. We measured oxygen consumption rates (OCR), glucose consumption, lactate production, ^13^C_6_-glucose flux, hypoxia inducible factor (HIF-1α) stability, intracellular lipid droplets after a lipid challenge, trans-epithelial electrical resistance, cell morphology, and pigmentation.

**Results:** Medium volumes commonly employed during RPE culture limit diffusion of O_2_ to cells, triggering hypoxia, activating HIF-1α, limiting OCR, and dramatically altering cell metabolism, with only minor effects on typical markers of RPE health. Media volume effects on O_2_ availability decrease acetyl-CoA utilization, increase glycolysis, and alter the size and number of intracellular lipid droplets under lipid-rich conditions.

**Conclusions:** Despite having little impact on visible and typical markers of RPE culture health, media volume dramatically affects RPE physiology “under the hood”. As RPE-centric diseases like age-related macular degeneration (AMD) involve oxidative metabolism, RPE cultures need to be optimized to study such diseases. We provide guidelines for optimal RPE culture volumes that balance ample nutrient availability from larger media volumes with adequate O_2_ availability seen with smaller media volumes.

## 1. Introduction

In the vertebrate eye, the retinal pigment epithelium (RPE) is a cell monolayer that supports photoreceptor health (1, 2, 3). The RPE is wedged between photoreceptors on its apical surface and a high-flow, fenestrated capillary bed called the choriocapillaris on its basolateral surface. The RPE is the primary site of degeneration in common blinding eye diseases, including Stargardt’s disease and age-related macular degeneration (AMD). Numerous lines of evidence support the importance of mitochondrial metabolism in RPE homeostasis: (i) the RPE is among the first tissues in the body affected in inherited mitochondrial disorders (4, 5), (ii) defects in RPE mitochondrial function result in proximal photoreceptor degeneration (6), and (iii) RPE degeneration in AMD is marked by significant mitochondrial structural abnormalities (7, 8). The RPE lives in a synergistic metabolic relationship with overlying photoreceptors, wherein photoreceptors produce lactate and succinate that the RPE consumes (9, 10). RPE mitochondria also consume lipids, which may be derived from phagocytosed photoreceptor outer segments or uptake of circulating lipoprotein particles (11, 12, 13).

Cells use energy from oxidation of fuels and reduction of O_2_ to drive ATP synthesis (14). In the terminal step of the mitochondrial electron transport chain, cytochrome C oxidase reduces O_2_ to H_2_O. Without O_2_, oxidation of metabolic intermediates upstream of cytochrome C oxidase is limited. Accumulation of reduced redox coenzymes under hypoxia leads to a metabolic shift towards glycolysis and lactate production, which generates energy less efficiently. To ensure an adequate O_2_ supply for mitochondria, nearly all animals have evolved circulatory and respiratory systems.

Culture of human primary and induced pluripotent stem cell (iPSC)-derived RPE is used to model RPE-based retinal degenerations (15, 16, 17, 18, 19, 20, 21). An important assumption in these studies is that atmospheric O_2_ concentrations do not limit mitochondrial activity. Estimates *in vivo* suggest that the fractional O_2_ availability at the RPE is ∼4-9% (extrapolated from (22)), whereas fractional atmospheric O_2_ is 21%, suggesting RPE *in vitro* could be in a hyperoxic environment. However, O_2_ dissolved in media and consumed by cultures must be replaced by diffusion from air above the medium. In accordance with Fick’s laws, O_2_ flux through the medium is constrained by the distance over which it must diffuse. The circulatory system delivers O_2_ continuously to within microns of a cellular target. In culture, O_2_ must diffuse across ∼3-25 mm of media, several orders of magnitude greater than diffusion distances *in vivo*. If cellular O_2_ consumption exceeds O_2_ diffusion through culture medium, cells will deplete the medium of O_2_ and lose the ability to oxidize fuels (23).

Herein, we test whether O_2_ consumption by RPE is greater than diffusion of O_2_ into the medium. We use multiple methods to show that widely used media volumes limit O_2_ availability in RPE cultures, stabilize the hypoxia sensor HIF-1α, increase glycolysis, impair mitochondrial metabolism, and alter intracellular lipid storage. Despite dramatic impacts on metabolism, other markers of RPE health are only subtly changed by prolonged exposure to higher media volumes. Lower media volumes restore O_2_ availability and mitochondrial metabolism but also result in faster nutrient depletion. We provide guidelines for how to balance media volumes and media change frequencies to preserve both nutrient availability and mitochondrial metabolism in cultured RPE.

## 2. Materials and Methods

### Cell Culture and Trans-Epithelial Electrical Resistance (TEER)

Primary human pre-natal RPE cultures (hfRPE) were grown and trans-epithelial electrical resistance (TEER) measured according to our detailed protocol outlined earlier (24, 25). All cultures demonstrated robust pigmentation, cobblestone morphology, and TEER of at least 500 Ω*cm^2^. All cells for metabolism analysis were passage 1, grown on porous 24-well inserts (Transwells; Corning; product 3470; cell growth area of 0.33 cm^2^) for at least 5 weeks before experimentation. Cells for Resipher experiments were grown on tissue-culture treated plastic in 96-well plates (Falcon; Corning; product 353072; cell growth area of 0.32 cm^2^) for at least 6 weeks prior to experimentation. Media compositions are based on standard RPE culture media (24, 25, 26, 27), with slight variations for each experiment, as outlined in **Supplemental Table 1**.

### Conjugation of Fatty Acids (FA) to Bovine Serum Albumin

Sodium palmitate (Sigma; product P9767) or sodium oleate (Sigma; product O7501) (FA) was solubilized at 70°C in 150 mM NaCl to create a 12.5 mM solution. The hot 12.5 mM FA solution was transferred to 1.7 mM FA-free BSA (MP Biomedical; product MP219989910) solubilized in glucose-free α-MEM (Supplementary Table 1) at a ratio of 0.8:1 and stirred at 37° C for 1 hour. The molar concentration of the conjugated FA-BSA solution was then adjusted by addition of 150 mM NaCl for a final concentration of 5 mM FA:0.85 mM BSA (6:1 molar ratio). Protocol was adapted from Seahorse Bioscience’s “Preparation of Bovine Serum Albumin (BSA)-Conjugated Palmitate.”

### HIF-1α Staining and Hypoxia Chambers

hfRPE on Transwells were placed in the incubator with either atmospheric O_2_ concentrations or in a hypoxia chamber (Billups-Rothenberg, Inc.; product MIC-101) in which O_2_ concentration was titrated to 8% utilizing a 1% O_2_ / 5% CO_2_ / 94% N_2_ gas mixture and an O_2_ sensor (Sensit P100 Personal Gas Leak Monitor) placed within the hypoxia chamber. After 24 hours of exposure, cells were lysed with 40 µL of 1.2X western blot SDS sample buffer. 100 µM of the prolyl hydroxylase inhibitor DMOG and 10 µM of the protease/phosphatase inhibitor MG-132 (Cell signaling; product 5872S) were used to prevent degradation of HIF. HIF-1α levels were determined in 25 µg lysate by standard SDS-PAGE and western blot techniques, utilizing a rabbit anti-HIF-1α antibody (Cell signaling; product 36169, 1:1000), a mouse monoclonal anti-GAPDH antibody (EnCor product MCA-1D4), and HRP-linked secondary antibodies (Goat anti-mouse HRP, Jackson Immuno, #115-035-062; Donkey anti-rabbit HRP, #711-035-152, Jackson Immuno).

### Lipid Loading and Lipid Droplet Staining

hfRPE on Transwells were loaded with 300 µM BSA-conjugated palmitate in serum-free media (Supplemental Table 1). Cells were incubated at 37°C for 18 hours and fixed with 4% paraformaldehyde / 4% sucrose solution for 15 minutes. After fixation, photochemical bleaching of the samples was performed. First, 150 μL of freshly made bleaching solution (50 µL deionized formamide, 50 µL of 3M sodium chloride/0.3M sodium citrate, 800 µL water, 163.4 µL 30% hydrogen peroxide) was added to the apical chamber of each Transwell. Next, the liquid light guide of an X-Cite 120Q microscopy system was centered 14 cm over the plate, held in place by a retort stand with utility clamp. After 2 minutes, Transwells were picked up and tapped gently to dislodge bubbles that form on the membrane surface. This step was repeated again at 5-7 minutes. Total bleach time was ∼15-20 minutes, but duration was slightly altered depending on the degree of pigment visible on the Transwell membrane towards the end of the bleaching process. Post-bleaching, Transwells were washed 5-8 times with 1x PBS to remove all bubbles. Cells were then quenched with a solution of 50 mM ammonium chloride in PBS for 10 minutes, permeabilized with 0.01% digitonin in PBS for 30 minutes at room temperature (RT), blocked with 3% bovine serum albumin (BSA) in PBS for 20 minutes, and incubated in primary antibody solution (anti-ADRP (perilipin-2), clone AP125 (Progen 610102), 1:10 dilution in 1% BSA in PBS) for 1 hour at RT or overnight at 4°C. Secondary antibody incubations and mounting were done in standard fashion, but avoiding all exposure to detergents. Hoechst-34580 (Sigma 63493, 1:500 dilution) was included in the secondary antibody incubation for 1 hour at RT.

### Microscopy

Z-stack images were obtained using the Leica STELLARIS 8 FALCON Confocal Microscope and image analysis was performed with Aivia 11 (Leica). To quantify LDs, a pixel classifier and meshes recipe was applied.

### Resipher

Resipher (Lucid Scientific, Atlanta, GA, USA) is an instrument that measures oxygen consumption rates (OCR) in standard multiwell culture plates. The device operates by measuring O_2_ concentrations across a range of heights in the media above the cell monolayer. These measurements correspond to the O_2_ gradient that forms in static media as oxygen diffuses through it. Using these gradient measurements, O_2_ flux through the media is calculated using Fick’s laws. An introduction to these laws and how they are used is available in the **Supplementary Discussion**.

Defined volumes of media were placed over cells and OCR was continuously monitored for up to 5 days without media change. Due to known uneven evaporative effects, wells on the edge of the plate were excluded from analysis. Each volume was tested in at least 6 replicate wells. All wells were confluent with a similar size and number of cells (**Supplemental Figure 1**).

### Glucose Concentration Assay

We measured media glucose with an enzymatic assay wherein glucose phosphorylation and oxidation is coupled to NADP^+^ reduction (28). NADPH absorbs light at 340 nm. We incubated 2-5 µL culture medium samples and 2-5 µL of 0-10 mM standards in the following assay buffer: 50 mM Tris, 1 mM MgCl_2_, 500 μM NADP^+^, 500 μM ATP, 0.2 U/mL Hexokinase, 0.08 U/mL glucose-6-phosphate dehydrogenase, pH 8.1. Using a BioTek Synergy 4 plate reader, we measured A_340_ over time at 37°C until it reached steady state. We used a linear fit of the standard curve to determine glucose concentrations. We determined glucose amount by multiplying concentrations with apical or basal medium volumes. A complete protocol for the glucose assay, including product numbers, is available at dx.doi.org/10.17504/protocols.io.dm6gpj5jdgzp/v1.

### Lactate Concentration Assay

We measured media lactate with an enzymatic assay where lactate dehydrogenase converts lactate and NAD_+_ to pyruvate and NADH (28). Pyruvate is consumed by the assay buffer, drawing the reaction to completion. NADH absorbs light at 340 nm. We incubated 2-5 µL culture medium samples and 2-5 µL of 0-20 mM standards in the following assay buffer: 300 mM glycine, 166 mM hydrazine, 2.5 mM NAD_+_, 8 U/mL lactate dehydrogenase. We determined A_340_ at 37°C with a Bio-Tek Synergy 4 plate-reader. Steady-state A_340_ values from standards were used to determine media [lactate]. A complete protocol for the lactate assay, including product numbers, is available at dx.doi.org/10.17504/protocols.io.6qpvr4733gmk/v1.

### Metabolite Extraction and Derivatization

Metabolites were extracted in 80% MeOH, 20% H_2_O supplemented with 10 µM methylsuccinate (Sigma; product M81209) as an internal standard. The extraction buffer was equilibrated on dry ice, and 150 µL was added to 2 µL of each medium sample. Samples were incubated on dry ice for 45 minutes to precipitate protein. Proteins were pelleted at 17,000 x g for 30 minutes at 4°C. The supernatant containing metabolites was lyophilized and stored at -80°C until derivatization.

Lyophilized samples were derivatized with 10 µL of 20 mg/mL methoxyamine HCl (Sigma; product 226904) dissolved in pyridine (Sigma; product 270970) and incubated at 37°C for 90 minutes. Samples were further derivatized with 10 µL tert-butyldimethylsilyl-N-methyltrifluoroacetamide (Sigma; product 394882) and incubated at 70°C for 60 minutes.

### Gas Chromatography-Mass Spectrometry

Metabolites were analyzed on an Agilent 7890/5975C GC-MS using methods described extensively in previous work (29). Briefly, 1 µL of derivatized sample is injected and delivered to an Agilent HP-5MS column by helium gas (1 mL/min). The temperature gradient starts at 100°C for 4 minutes and increases 5°C/min to 300°C, where it is held for 5 minutes. We used select ion monitoring (SIM) to record ions (m/z: ∼50–600) in expected retention time windows. Peaks were integrated in MSD ChemStation (Agilent). We corrected for natural isotope abundance using IsoCor (30). Corrected metabolite signals were converted to molar amounts by comparing metabolite peak abundances in samples with those in a standard mix. Multiple concentrations of this mix were extracted, derivatized, and run alongside samples in each experiment.

### Statistical Analysis

All figures representing data in this manuscript represent either individual data points or the arithmetic mean, ± standard error. Western blot data was analyzed with one-way ANOVA or a student’s t-test. The significance threshold for statistical tests was set as: *p≤0.05, ***p≤0.001. All statistics were calculated using Prism v9.5.0 (GraphPad Software, LLC)

## 3. Results

### 3.1. The volume of cell culture media determines the balance between hypoxic conditions and nutrient availability, with only subtle effects on markers of RPE health

Primary human RPE (hfRPE) cultures were grown on Transwell filters in 24-well plates to best mimic cellular polarity and access to nutrients *in vivo*. We have previously established these cultures as highly polarized and differentiated, with high TEER, expression of RPE-specific expression markers, high pigmentation, and capacity for photoreceptor outer segment (OS) phagocytosis (24). The apical side of the filter faces the air-medium interface, and the basal surface faces the bottom of the culture plate.

O_2_ can reach cells through either chamber (**Figure 1A**) but the apical chamber media directly touches cells without an intervening membrane and unrestricted access to the atmosphere, so we chose to alter apical media volumes to understand the effect of media volume on O_2_ availability at the cell monolayer. We hypothesized that O_2_ availability and therefore O_2_ consumption rate (OCR) are limited by medium depth under standard culture conditions, where apical medium volume can vary between 100 µL to 200 µL or more. To test this hypothesis, we grew hfRPE on 96-well plastic plates, which have the same surface area as a 24-well Transwell. We measured OCR in wells with 65, 95, 125, or 200 µL of media. The Resipher instrument determines OCR based on the vertical [O_2_]-gradient that develops within the medium in the well. OCR is lowest at 200 µL (<100 fmol/mm^2^/second) but increases with lower apical medium volumes (**Figure 1B,C**), showing that OCR is indeed limited by medium volume. Steady-state OCR is nearly identical at 65 µL and 95 µL medium depth (140-160 fmol/mm^2^/second), suggesting that at volumes just under 100 µL, [O_2_] no longer limits oxidative phosphorylation (**Figure 1B,C**). A similar relationship between medium volume and OCR rates was confirmed in human iPSC-RPE cultures. Given a cell culture surface area of 0.33 cm^2^, this corresponds to a media volume to surface area ratio of 300 µL/cm^2^. These OCR values, stable for almost a day, represent how the cultures initially respond to O_2_ availability. With additional time, cells situated under larger medium volumes further adapt their OCR to limited O_2_ availability.

Our data imply that cultured human RPE could be hypoxic when grown at >300 µL/cm^2^. To confirm that cells under higher media volumes were sensing hypoxia, we measured levels of the hypoxia responsive transcription factor, HIF-1α. HIF-1α is degraded when hydroxylated, and the ability of cells to hydroxylate HIF-1α depends on the availability of O_2_ (31). Thus, low O_2_ concentrations increase HIF-1α levels. We treated hfRPE on Transwells at 8% atmospheric or 21% atmospheric O_2_ for 24 hours, and at each atmospheric O_2_ concentration also modulated apical medium volume, while maintaining basolateral media volume constant at 400 µL. Western blot of hfRPE lysate revealed a clear effect of both medium depth and of atmospheric [O_2_] on HIF-1α, supporting the conclusion that increasing medium volumes can lead to hypoxia in culture (**Figure 1D**). Notably, while 8% O_2_ is considered physioxic *in vivo*, 8% atmospheric O_2_ increases HIF-1α levels *in vitro*. Even under the lowest culture volume, 65 µL, HIF-1α was significantly more stabilized in the 8% O_2_ conditions than the 21% O_2_ conditions (**Figure 1D;** bottom right). This reaffirms that high diffusion distance *in vitro* limits O_2_ access to cells and complicates comparisons of O_2_ concentrations *in vivo* with O_2_ concentrations *in vitro*.

**Figure 1.**
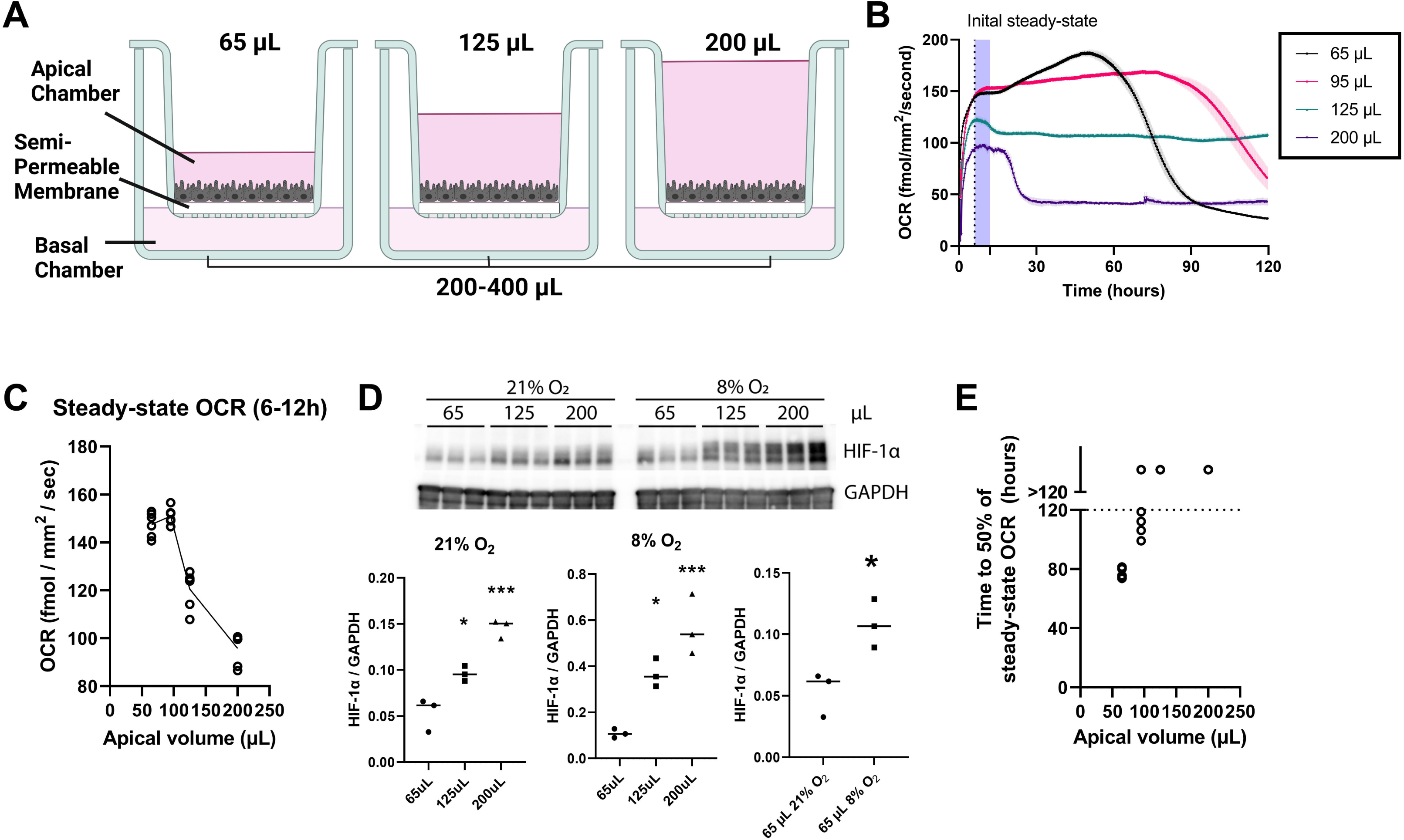
Medium volume limits cellular O_2_ uptake by increasing O_2_ diffusion distance. **(A)** Schematic showing the path O_2_ must take to get from air, through liquid barriers (culture medium), and to cells at varying apical volumes and constant basal volume. (**B**) OCR over time for wells at different medium depths (65, 95, 125, 200 µL; n=6 wells/group). (**C**) Average initial steady-state OCR at each medium depth. (**D**) [O_2_]-dependent or medium volume-dependent changes in HIF-1α, as assessed by Western blot (normalized to GAPDH) (n=3). (**E**) Medium volume also affects mitochondrial fuel availability. As these fuels are depleted, OCR drops. Plotted is the time (in hours) after media change until OCR drops to 50% of its previous steady-state (n=6). *p≤0.05, ***p<0.001, one-way ANOVA or student’s T-test.

While lower media volumes improve O_2_ availability, they also limit nutrient supply. The typical interval between medium changes in RPE cultures is 48-72 hours. To measure the effects of media volume on substrate depletion rates, we supplied 65, 95, 125, or 200 µL medium to hfRPE and measured OCR up to 120 hours. When all mitochondrial substrates have been consumed, OCR will drop. With 65 µL, the time to a 50% drop from steady-state OCR is 76.7 ± 1.3 (mean ± SEM) hours after a medium change. With 95 µL, OCR dropped to 50% by 109 ± 4.2 hours after a medium change. At greater medium depths, OCR persisted at steady-state levels past 120 hours. Thus, at media volumes just under a ratio of 300 µL/cm^2^, RPE grows without O_2_ limitation and requires media changes at least twice weekly to prevent a decrease in OCR from insufficient fuel availability (**Figure 1E**).

To determine the effects of media volume on typical markers of RPE health, we examined tight-junction integrity, morphology, and pigmentation (32). We previously established trans-epithelial electrical resistance (TEER) as a sensitive marker for subtle cell death (33). hfRPE cultures supplied with 100 µL or 200 µL of media for up to three weeks had no differences in cell morphology or pigmentation (**Supplemental Figure 2A**), but did have subtle differences in TEER that were present only after exposure for 3 weeks and not 1 week (**Supplemental Figure 2B**). These results suggest slow dysfunction in high media-volume cultures over time that are not apparent by visual inspection.

### 3.2. Limiting O2-dependent metabolism accelerates glycolysis, depleting glucose faster and causing an accumulation of lactate

When O_2_-availability limits mitochondrial metabolism, cells can process glucose by reducing pyruvate into lactate. This mode of glucose utilization produces ATP at ∼1/16^th^ the efficiency as complete oxidation of glucose to CO_2_ by fully coupled mitochondria, so more glucose is consumed to support the same energetic needs.

As a proof of this concept, we examined the effects of limiting O_2_ availability on glucose consumption and lactate production in hfRPE cultures. We cultured cells on Transwells with 100 µL apical medium and 200 µL basolateral media over 24 hours, where the cells were equilibrated with 21% O_2_ in air. Next, we incubated the same cells in the same media volume but set O_2_ at 8% in air. We measured total glucose and total lactate content in culture medium before it was exposed to cells and after 24 hours with an enzymatic assay. 8% O_2_ increases glucose utilization (**Figure 2A**) and lactate production (**Figure 2B**) such that almost all glucose provided to RPE cells was consumed by 24 hours.

**Figure 2.**
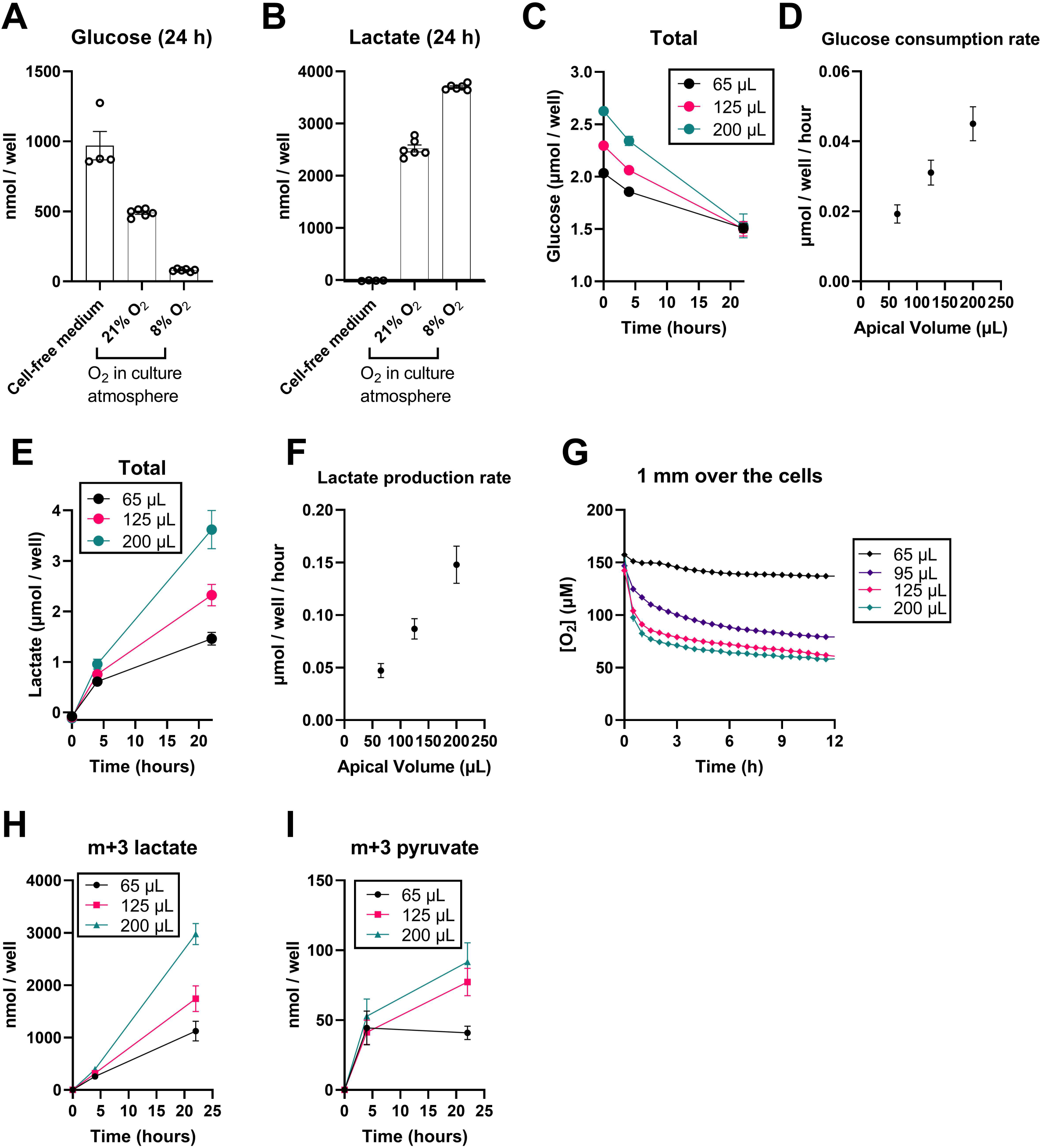
Glycolysis is accelerated by increasing medium depth. **(A)** Medium glucose and **(B)** medium lactate amounts after 24 hours of culture in medium without cells, medium with cells at 21% atmospheric O_2_, or cells at 8% atmospheric O_2_. Decreased O_2_ levels increase glucose utilization and lactate production. **(C-D)** Increasing medium volume also limits cellular O_2_ availability, increasing glucose consumption and **(E-F)** lactate production. **(G)** Limited cellular O_2_ availability develops only after several hours in culture as steady-state O_2_ diffusion gradients develop. This explains why medium volume effects on glucose and lactate (C-F) develop only after 4 hours. **(H-I)** Decreased glucose levels and increased lactate levels are a direct consequence of glucose consumption, as assessed by ^13^C_6_-glucose tracing to m+3 lactate **(H)** and m+3 pyruvate **(I)** (n=5-6).

To determine whether higher medium volumes have the same effect on glucose consumption and lactate production as 8% O_2_, we cultured hfRPE in ambient O_2_ (21%) for 22 hours in 65, 125, or 200 µL of apical media and 400 µL of media basolaterally. We replaced normal unlabeled glucose (5 mM) with the same concentration of ^13^C_6_-glucose and supplemented medium with 150 µM palmitate-BSA and 150 µM oleate-BSA to provide substrates used by mitochondria. We probed metabolite release into the apical and basal chambers through a combination of mass spectrometry and enzymatic assays. The metabolite amounts represent the sum of apical and basal metabolite content (**Figures 2, 3**).

**Figure 3.**
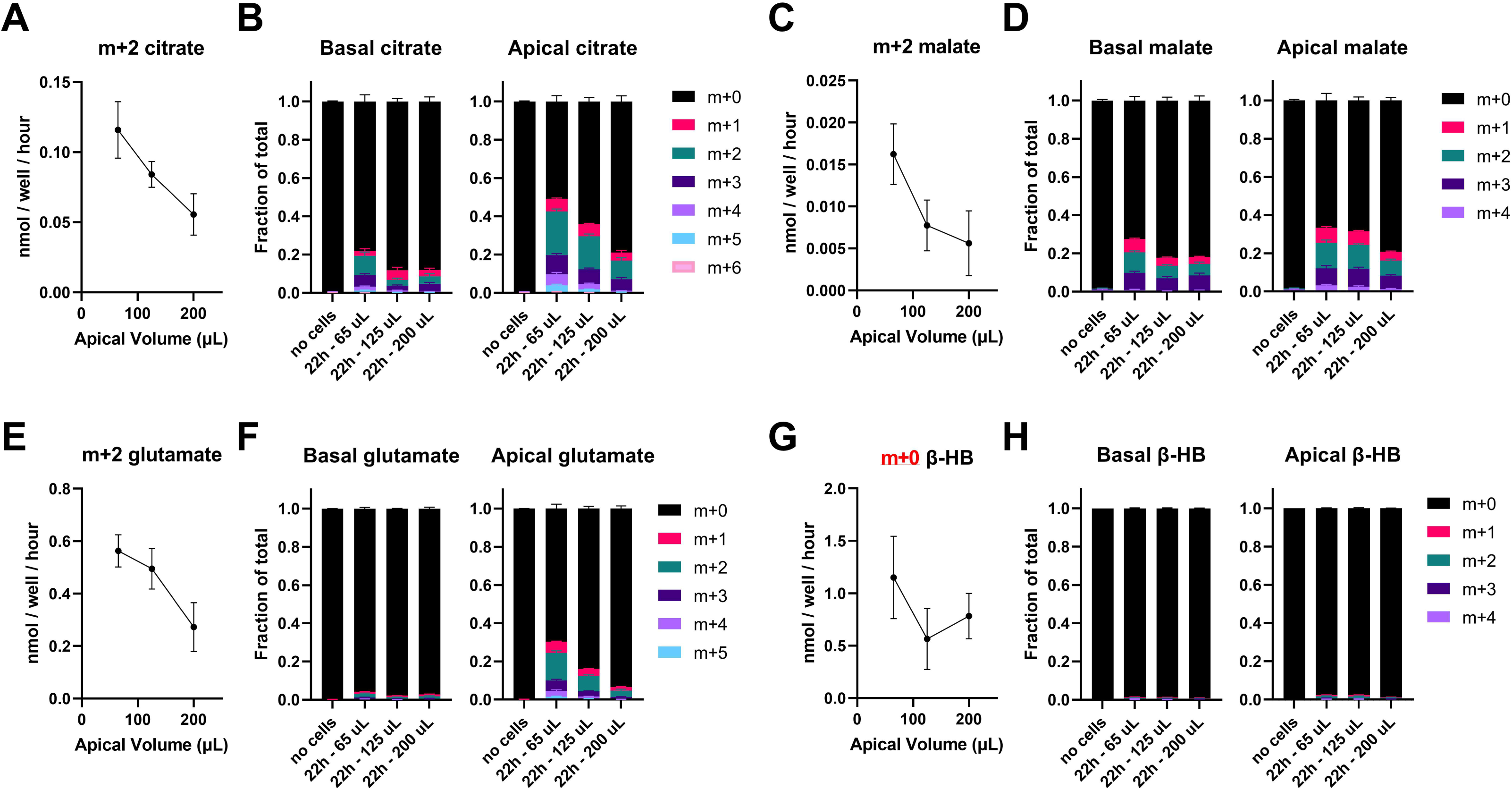
Mitochondrially-dependent acetyl-CoA utilization increases with decreasing medium depth. We traced the fate of ^13^C_6_-glucose into mitochondrially-dependent intermediates, measuring the rate of appearance of m+2 labeled intermediates in the apical media from 4-22 hours after media change (**A**,**C**,**E)**, as well as the fraction of each intermediate in the apical vs. basolateral media that picked up a ^13^C label at 22 hours **(B**,**D**,**F)**. The apical media secretion rate of citrate **(A)**, malate **(C)**, and **(E)** glutamate derived from glucose (m+2) decreases with higher media volumes. In addition, the fraction of total **(B)** citrate, **(D)** malate, and **(F)** glutamate with a ^13^C label decreases at higher media volumes. These results confirm that limited O_2_ diffusion to cells prevents ongoing mitochondrial activity. **(G)** The total and unlabeled (m+0) β-HB in the apical media were almost identical, suggesting that glucose does not contribute to ketogenesis. β-HB present in apical media likely derives from unlabeled fatty acids added to the cultures. Similar to other mitochondrially-dependent intermediates, β-HB production was highest at the lowest media volume where O_2_ availability is not limiting. **(H)** Fractional labeling of β-HB from ^13^C_6_-glucose confirms almost no production of β-HB from glucose (n=5-6).

Like when cells are cultured at a low O_2_ concentration, RPE cell glucose consumption (**Figure 2C-D**) and lactate production (**Figure 2E-F**) from 4-22 hours increases with greater apical medium depth. Similar results relating medium depth to glucose consumption and lactate production were found in human iPSC-RPE cultures. For the first 4 hours after a medium change, glycolytic flux is insensitive to apical medium depth, and differences between conditions only appear between 4 and 22 hours in culture. This latency required to detect changes in glycolysis occurs because O_2_ gradients take time to develop, and flux differs only after steady-state O_2_ gradients form. We confirmed this by measuring O_2_ concentration 1 mm above the cell monolayer and found that O_2_ availability became progressively more limited after several hours in wells with higher apical medium volumes (**Figure 2G**).

Pyruvate and other fuels in culture medium could be sources of lactate production. To follow carbons from glucose specifically, we quantified m+3 glycolytic products, derived from ^13^C_6_-glucose. After a steady-state O_2_ gradient is established in culture (4 hrs), m+3 lactate (**Figure 2H**) and m+3 pyruvate (**Figure 2I**) both increase in an apical volume-dependent manner. This indicates an acceleration in glycolysis as medium depth increases.

### 3.3. Acetyl-CoA utilization increases with decreasing medium depth

At lower apical medium volumes, more O_2_ diffuses to mitochondria, which should facilitate mitochondrial TCA cycle activity and β-oxidation. Carbons from ^13^C_6_-glucose are used in glycolysis to make m+3 pyruvate, which is then decarboxylated into m+2 acetyl-CoA. m+2 acetyl-CoA enters the TCA cycle and makes m+2 citrate and downstream m+2 intermediates. Labeled intermediates produced from acetyl-CoA are exported from cells into medium, so sampling media is a noninvasive way to assess metabolism of ^13^C tracers.

Growing hfRPE on Transwells under conditions identical to those in Figure 2, we found a clear relationship between media depth and labeled intermediates released into media. As apical medium depth decreases, more m+2 citrate is made (**Figure 3A**) and a greater proportion of citrate is ^13^C-labeled (**Figure 3B**). The same trend exists for downstream TCA cycle and anapleurotic intermediates such as malate (**Figure 3C-D**) and glutamate (**Figure 3E-F**).

β-hydroxybutyrate (β-HB) is a ketone body made using carbons from two acetyl-CoA molecules, and the RPE’s capacity for ketogenesis is well established (12, 15). While glucose can fuel β-HB production, ketogenesis is considered primarily to be a product of fatty acid metabolism (34, 35) that occurs in the mitochondria. Compared with exported TCA cycle intermediates, most β-HB produced by hfRPE over 22 hours is unlabeled (m+0) (**Figure 3G-H**), suggesting that it is not derived from ^13^C_6_-glucose but instead from unlabeled fatty acids that we provided. As with TCA cycle intermediates, production of m+0 β-HB also increased at the lowest medium volume (**Figure 3G**), suggesting fatty acid metabolism in RPE culture may be limited at higher medium volumes.

### 3.4. Mass action is not the sole driver of differences in metabolite release rates between different volume conditions

The law of mass action states that reaction rate is proportional to concentration of products and reactants. Applied to metabolite export into medium, the law suggests that the ratio of the metabolite’s intracellular and extracellular concentration will determine the rate of export. Higher medium volumes dilute metabolites more, and thus facilitate faster export. Thus, medium depth can affect cellular metabolism not only by altering cellular O_2_ and nutrient availability, but also through mass action. We tested whether this variable confounds the changes in secreted metabolites we observe from differences in O_2_ supply.

While apical volumes in our experimental set-up differ, basal volumes are constant. Thus, metabolite efflux to the basal side should exclusively reflect O_2_-dependent differences in hfRPE metabolism. When labeling of lactate, pyruvate, and citrate from ^13^C_6_-glucose is measured separately in apical and basal chambers, there is a clear effect of volume on the flux of m+3 lactate (**Figure 4A**), m+3 pyruvate (**Figure 4B**), and m+2 citrate (**Figure 4C**) in both chambers. Thus, while mass action may play a general role in metabolite release, medium depth-dependent differences in O_2_ drive differential metabolite release to the basal chamber.

**Figure 4.**
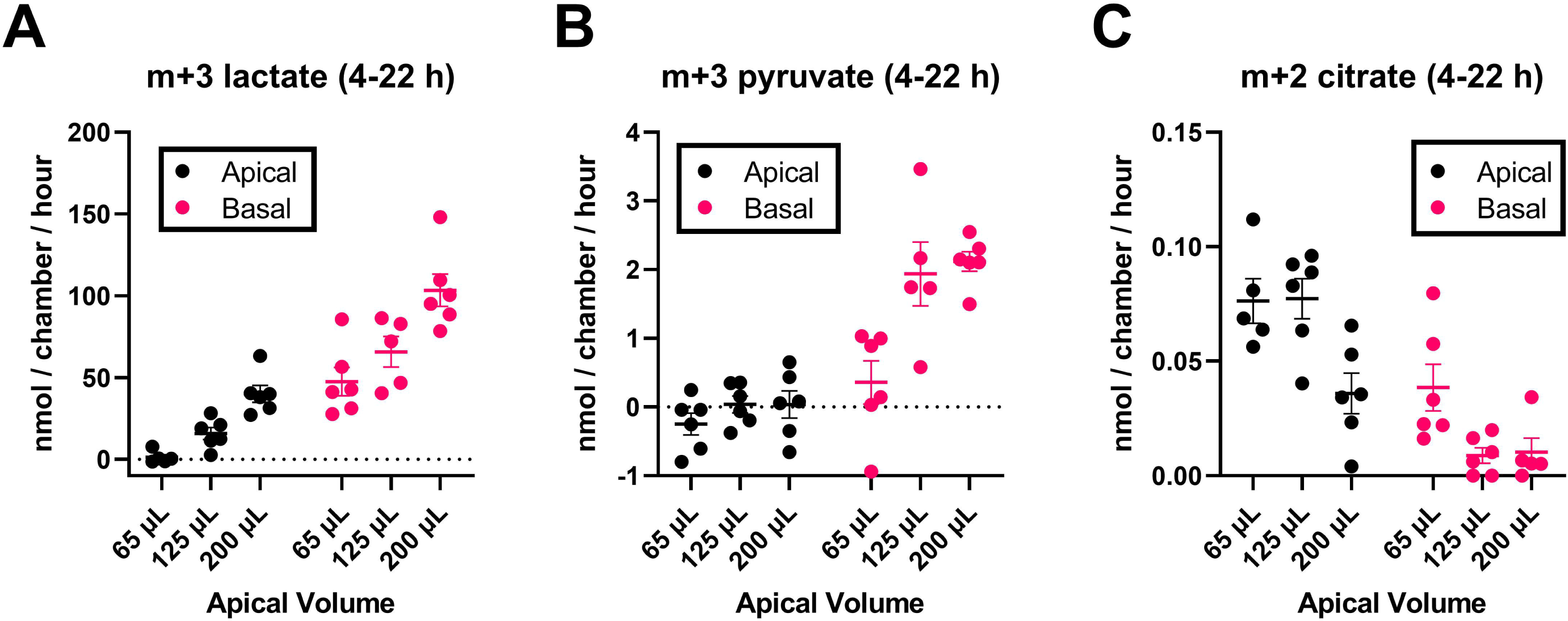
Volume effects on metabolite release rates depend largely on O_2_ availability. Given different apical volumes, differences in metabolite release rates into the medium could be caused by differences in O_2_ availability or simply mass action effects. While apical media volumes differ, basal media volumes are constant across all conditions. Thus, release of metabolites into the basolateral media will be unaffected by mass action, and any apical media volume effects on basal metabolite release must be due to differences in O_2_ availability between the conditions. **(A)** The release rate of m+3 lactate derived from ^13^C_6_-glucose into the apical (black dots) or basal (pink dots) chambers from 4-22 hours are similarly affected by apical media volume, confirming O_2_ availability is the dominant factor affecting release. Similar analysis for **(B)** pyruvate and **(C)** citrate.

### 3.5. Media volume affects lipid droplet dynamics after RPE lipid challenge

The RPE is a prolific consumer of lipids (36) and forms temporary lipid droplets (LDs) in response to high lipid concentrations (37). The ability of cells to metabolize fatty acids from lipid droplets partly controls the size and persistence of LDs (38). β-oxidation of fatty acids is O_2_-dependent, so we hypothesized that higher media volumes would limit β-oxidation and lead to an accumulation of LDs. To test this hypothesis, we fed hfRPE serum-free media supplemented with 300 µM BSA-conjugated palmitate for 18 hours, comparing LD formation with an apical volume of 65 µL vs. 200 µL (with a constant basal volume of 400 µL and 21% atmospheric O2). As a control for limited O_2_ availability but equal absolute amounts of palmitate, we also provided cells with 65 µL of medium under 1% atmospheric O_2_. Decreasing O_2_ levels via 1% atmospheric O_2_ or higher media volumes had a similar effect on increasing LD amount and size (**Figure 5A-C**). Interestingly, larger LD were also more likely to be localized basolaterally (**Figure 5D**). Similar effects of medium volume on LD dynamics were found using iPSC-RPE. Thus, studies exploring the role of lipid metabolism and LDs in the RPE need to be particularly cognizant of media volume effects.

**Figure 5.**
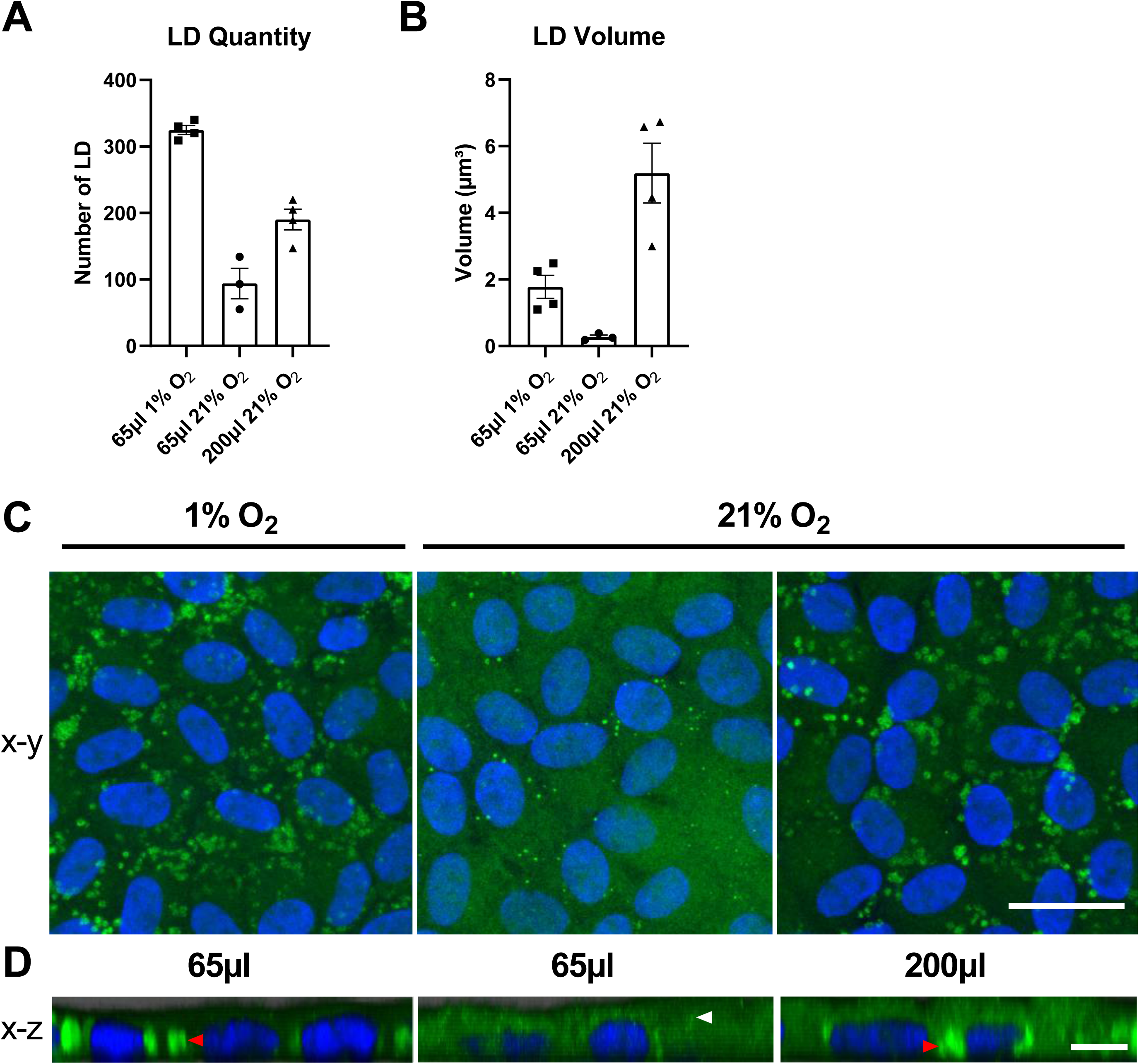
Media volume affects RPE lipid droplet size, number, and localization after lipid challenge. In both hypoxic (1% O_2_) and high media volume (200 µL) conditions, there is an increase in the **(A)** number of lipid droplets (LD) and **(B)** size of LD formed after 18 hour lipid challenge (300 µM palmitate). n=3-4 randomly selected images. **(C)** Immunostaining LD (ADRP/perilipin-2, green) and nuclei (Hoechst 34580, blue) under high volume or 1% O_2_ conditions. **(D)** The larger LD that form under high volume or low atmospheric O_2_ conditions are predominantly basolateral (red arrowhead), adjacent to nuclei. The smaller LD that form under normoxia are more apically localized (white arrowhead). Scale bar= 10 µM for X-Y and X-Z images.

## 4. Discussion

While primary and iPSC-derived RPE culture models have shown great promise in recapitulating characteristics of RPE in health and disease, we show that media volumes commonly employed for RPE culture can profoundly limit O_2_ availability. This impacts everything from mitochondrial metabolism to activation of hypoxia-responsive transcription factors to lipid dynamics. These data mirror findings on the impact of media volume in adipocytes (23). Despite profound effects of media volume on RPE central carbon and lipid metabolism, we noted little change in commonly employed markers for RPE health. There were no effects on RPE morphology and pigmentation, and only minor effects on TEER after prolonged (3 week) exposure to high media volumes. This uncoupling between highly altered cellular metabolism and unaltered RPE function/survival could explain why effects of media volume on RPE cultures have largely been ignored.

Despite the lack of overt effects of medium depth on many markers of RPE culture health, it is still essential that RPE metabolism be modeled accurately. Hypoxia inducible factor 1α (HIF-1α) is stabilized in media volumes commonly employed for RPE culture, yet given HIF’s documented role in various RPE pathologies (39, 40, 41), investigating RPE biology under conditions that constitutively activate HIF could be misleading. Numerous studies have established that LDs increase dramatically under hypoxia or conditions that activate HIF (40, 42). In our cultures, limited O_2_ availability from higher media volumes altered LD number, size, and polarized intracellular localization after lipid challenge. Interestingly, a prior paper demonstrated that hypoxia is known to specifically stimulate larger LDs (43). Similarly, the RPE pathogenesis of AMD is thought to involve RPE lipid handling (44). There is intense interest in understanding and manipulating RPE β-oxidation for therapeutic purposes in this disease. Our data suggests that *in vitro* studies of lipid metabolism could be hampered by limited O_2_ availability.

Cellular O_2_ availability depends not just on media depth, but also on the cell monolayer’s OCR. When OCR is higher, O_2_ is depleted faster and lower media depths are necessary to ensure adequate O_2_ supply. In this study, hfRPE OCR necessitates a volume/surface area ratio of 300 µL/cm^2^ or lower to avoid hypoxia at the cell monolayer. Media volume to surface area ratios lower than 300 µL/cm^2^ would increase O_2_ availability but are not necessary because they do not further increase OCR and could restrict nutrient availability. At a volume/surface area ratio of approximately 200 µL/cm^2^ (65 µL in 24-well Transwell), nutrient depletion *begins* to affect mitochondrial metabolism after approximately 50-55 hours, whereas at a ratio of approximately 300 µL/cm^2^ (95 µL in 24-well Transwell), nutrient depletion doesn’t *begin* to affect mitochondrial metabolism until approximately 80-85 hours (**Figure 1B**). Since most labs change cell culture media between 2-3 times per week, a volume/surface area ratio of approximately 300 µL/cm^2^ balances O_2_ supply and nutrient availability for hfRPE.

Factors other than media depth and OCR also influence O_2_ availability *in vitro*. These include diffusion of O_2_ through the plastic sides and bottom surface of the well, humidity in the incubator, and the atmospheric pressure (e.g. - in labs at elevation) (45). These effects can only partially be controlled but can be compensated for by altering medium volume. To estimate the consequences of manipulating medium volume and other parameters on cellular O_2_ availability, we designed an interactive web notebook, available at: https://observablehq.com/@lucid/oxygen-diffusion-and-flux-in-cell-culture. The assumptions and calculations underpinning this calculator are outlined in the **Supplementary Discussion**.

In conclusion, media depth is a critical factor in determining O_2_ availability in RPE culture, with myriad serious consequences disguised underneath normal-appearing morphology and pigmentation. Unfortunately, medium volume is rarely reported in studies on RPE culture (45), including our own prior publications. To ensure more consistency across studies on RPE metabolism and mitochondrial health, we recommend reporting cell culture surface area, confluency, media volume, and time between media change and the end-point of the experiment. For RPE cultures with OCR rates similar to hfRPE, we recommend maintaining cultures at 300 µL of media per cm_2_ of surface area with no more than four days between media changes.

## Supporting information

Supplemental Discussion and Supplemental Figure 3

## Abbreviations

O2: molecular oxygen
OCR: oxygen consumption rate
RPE: retinal pigment epithelium
iPSC-RPE: induced pluripotent stem cell derived retinal pigment epithelium
hfRPE: human pre-natal retinal pigment epithelium
LD: Lipid droplet

## Acknowledgements

Figure panel 1A was created with BioRender.com. We thank Dr. Subramanian Pennathur for helpful comments.

## Figure captions

**Supplemental Figure 1. Number of highly differentiated confluent RPE cells is consistent between cell culture wells.** Manually counted cell numbers from same size images randomly taken in the center of 8 wells were normalized to the average.

**Supplemental Figure 2. Higher media volumes cause mild alterations in typical markers of RPE health. (A)** Brightfield images show similar polygonal shape and pigmentation for RPE under 100 µL and 200 µL apical (basal 600 µL) media volume after 3 weeks. Scale bar = 10 µM. **(B)** TEER was measured at day 0, 7, and 21 days after culturing in 100 µL or 200 µL apical media (basal 600 µL), with subtle differences emerging only after 3 weeks. ns, not significant, *p≤0.05. TEER readings were normalized to starting TEER value on day 0 and 100 µL value. n=6.

