## Supplemental Table 1 for "Medium depth influences O_2_ availability and metabolism in cultured RPE cells"

| Experiment | Figure 2C-F, H-I<br>Figure 3<br>Figure 4 | Figure 1B,C,E<br>Figure 2G<br>Supplemental Figure 2 | Figure 1D<br>Figure 2A-B | Figure 5 |
|---|---|---|---|---|
| aMEM (Alpha Modification of Eagle's Media) + Non-Essential Amino Acids Solution | USBiological Life Sciences<br>M3852-02<br>+<br>Gibco 11140050 | Corning<br>15-012-CV<br>+<br>Gibco 11140050 | Corning<br>15-012-CV<br>+<br>Gibco 11140050 | Corning<br>15-012-CV<br>+<br>Gibco 11140050 |
| *Inorganic Salts* | | | | |
| *Units* | mg/L | mg/L | mg/L | mg/L |
| CaCl2 (anhydrous) | 265.00 | 200.00 | 200.00 | 200.00 |
| KCl | 400.00 | 400.00 | 400.00 | 400.00 |
| MgSO4 (anhydrous) | 97.67 | 97.70 | 97.70 | 97.70 |
| NaCl | 6800.00 | 6800.00 | 6800.00 | 6800.00 |
| NaH2PO4 • H2O | 122.00 | 140.00 | 140.00 | 140.00 |
| NaHCO3 | 2200<br>(supplemented in, does not come with product originally) | 2200.00 | 2200.00 | 2200.00 |
| *Amino Acids* | | | | |
| *Units* | mg/L | mg/L | mg/L | mg/L |
| L-Alanine | 33.90 | 33.90 | 33.90 | 33.90 |
| L-Arginine • HCl | 126.00 | 126.40 | 126.40 | 126.40 |
| L-Asparagine • H2O | 63.20 | 63.20 | 63.20 | 63.20 |
| L-Aspartic acid | 43.30 | 43.30 | 43.30 | 43.30 |
| L-Cysteine • HCl • H2O | 100.00 | 100.00 | 100.00 | 100.00 |
| L-Cystine • 2HCl | 31.30 | 31.20 | 31.20 | 31.20 |
| L-Glutamic acid | 89.70 | 89.70 | 89.70 | 89.70 |
| L-Glutamine | 292.00 | -- | -- | -- |
| L-Alanyl-L-Glutamine (GlutaMAX) | -- | 445.00 | 445.00 | 445.00 |
| Glycine | 57.50 | 57.50 | 57.50 | 57.50 |
| L-Histidine • HCl • H2O | 42.00 | 41.90 | 41.90 | 41.90 |

| | | | | |
|---|---|---|---|---|
| L-Isoleucine | 52.00 | 52.50 | 52.50 | 52.50 |
| L-Leucine | 52.00 | 52.50 | 52.50 | 52.50 |
| L-Lysine • HCl | 72.50 | 72.50 | 72.50 | 72.50 |
| L-Methionine | 15.00 | 15.00 | 15.00 | 15.00 |
| L-Phenylalanine | 32.00 | 32.50 | 32.50 | 32.50 |
| L-Proline | 51.50 | 51.50 | 51.50 | 51.50 |
| L-Serine | 35.50 | 35.50 | 35.50 | 35.50 |
| L-Threonine | 48.00 | 47.60 | 47.60 | 47.60 |
| L-Tryptophan | 10.00 | 10.00 | 10.00 | 10.00 |
| L-Tyrosine • 2Na | 52.00 | -- | -- | -- |
| L-Tyrosine • 2Na • 2H2O | -- | 51.90 | 51.90 | 51.90 |
| L-Valine | 46.00 | 46.80 | 46.80 | 46.80 |
| *Vitamins* | | | | |
| *Units* | mg/L | mg/L | mg/L | mg/L |
| Ascorbic acid | 50.00 | 50.00 | 50.00 | 50.00 |
| Biotin | 0.10 | 0.10 | 0.10 | 0.10 |
| D-Calcium pantothenate | 1.00 | 1.00 | 1.00 | 1.00 |
| Choline chloride | 1.00 | 1.00 | 1.00 | 1.00 |
| Folic acid | 2.00 | 1.00 | 1.00 | 1.00 |
| i-Inositol | 1.00 | 2.00 | 2.00 | 2.00 |
| Nicotinamide | 1.00 | 1.00 | 1.00 | 1.00 |
| Pyridoxine • HCl | 1.00 | 1.00 | 1.00 | 1.00 |
| Riboflavin | 0.10 | 0.10 | 0.10 | 0.10 |
| Thiamine • HCl | 1.00 | 1.00 | 1.00 | 1.00 |
| Vitamin B12 | 1.36 | 1.36 | 1.36 | 1.36 |
| *Other* | | | | |
| *Units* | mg/L | mg/L | mg/L | mg/L |
| Adenosine | -- | -- | -- | -- |
| Cytidine | -- | -- | -- | -- |
| 2’-Deoxyadenosine • H2O | -- | -- | -- | -- |

| | | | | |
|---|---|---|---|---|
| 2'-Deoxycytidine • HCl | -- | -- | -- | -- |
| 2'-Deoxyguanosine • H2O | -- | -- | -- | -- |
| D-Glucose | glucose) | 1000.00 | 1000.00 | 1000.00 |
| Guanosine | 10.00 | -- | -- | -- |
| Lipoic Acid | 0.20 | 0.20 | 0.20 | 0.20 |
| Phenol Red | 11.00 | 10.00 | 10.00 | 10.00 |
| Sodium Pyruvate | (supplemented in, does | 110.00 | 110.00 | 110.00 |
| Thymidine | -- | -- | -- | -- |
| Uridine | -- | -- | -- | -- |
| *N1 Media Supplement* | Sigma N6530 | Sigma N6530 | Sigma N6530 | Sigma N6530 |
| *Units* | µg/mL | µg/mL | µg/mL | µg/mL |
| Recombinant human insulin | 5.0000 | 5.0000 | 5.0000 | 5.0000 |
| Human transferrin (partially iron-saturated) | 5.0000 | 5.0000 | 5.0000 | 5.0000 |
| Sodium selenite | 0.0050 | 0.0050 | 0.0050 | 0.0050 |
| Putrescine | 16.0000 | 16.0000 | 16.0000 | 16.0000 |
| Progesterone | 0.0073 | 0.0073 | 0.0073 | 0.0073 |
| *Additions To Culture Media* | | | | |
| *Units* | mg/L | mg/L | mg/L | mg/L |
| U-13C-D-Glucose (CLM-1396) | 1000 | -- | -- | -- |
| *Units* | U/mL | U/mL | U/mL | U/mL |
| Penicillin-Streptomycin (Gibco 15140-122) | 100 | 100 | 100 | 100 |
| *Units* | mg/L | mg/L | mg/L | mg/L |
| Taurine (Sigma T8691) | 250.00 | 250.00 | 250.00 | 250.00 |
| *Units* | µg/L | µg/L | µg/L | µg/L |
| Hydrocortisone-Cyclodextrin (Sigma H0396) | 20 | 20 | 20 | 20 |
| (Sigma T5516) | 0.013 | 0.013 | 0.013 | 0.013 |
| *Concentration* | mM | mM | mM | mM |
| L-carnitine (Thermo A1761809) | 400 | -- | -- | -- |
| BSA conjugated-Palmitate | 150 | -- | 150 | 300 |

| | | | | |
|---|---|---|---|---|
| BSA conjugated-Oleate | 150 | -- | 150 | -- |
| *Percentage* | % | % | % | % |
| Fetal Bovine Serum (Bio-Techne S11550H) | 0 | 5 | 0 | 0 |
