## Supplementary figures and images for "Medium depth influences O_2_ availability and metabolism in cultured RPE cells"

### Supplemental Figure 1

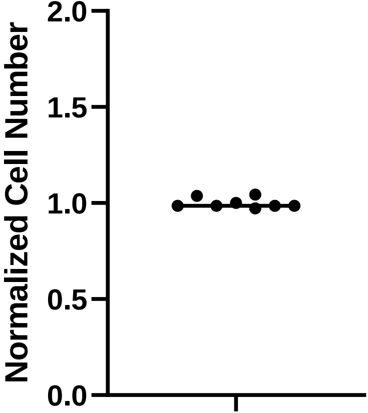

### Supplemental Figure 2

A

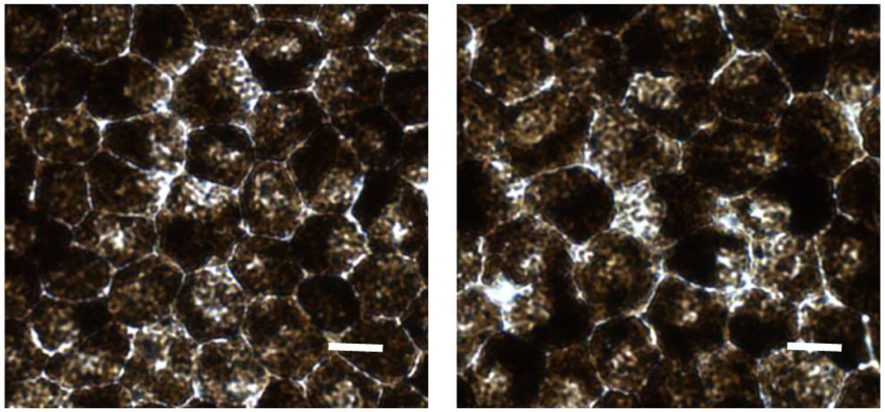

100 µL 200 µL

B

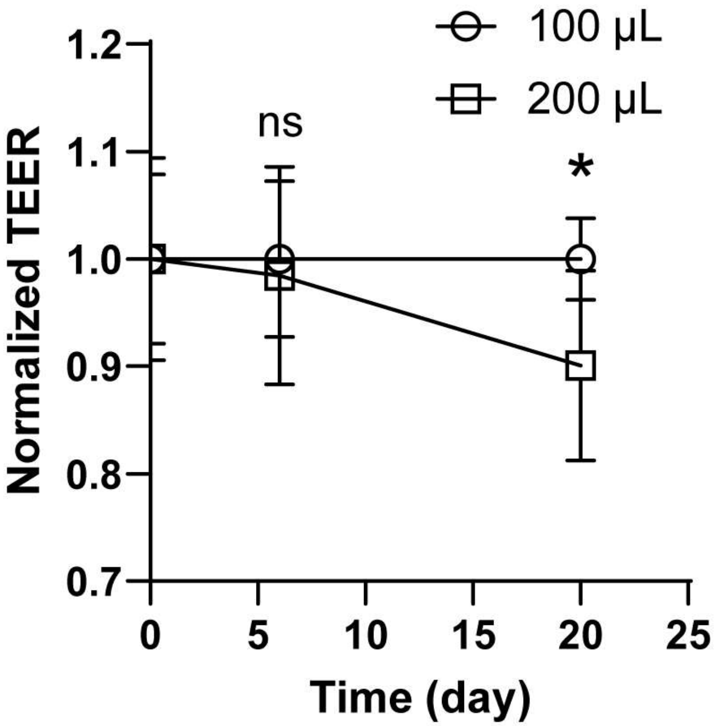
