## Supplemental Discussion and Supplemental Figure 3 for "Medium depth influences O_2_ availability and metabolism in cultured RPE cells"

The purpose of this discussion is to show the calculations and assumptions necessary for estimating $O_2$ levels available to cells in culture, and the culture parameters that affect $O_2$ availability. This discussion is accompanied by an online calculator that allows the reader to adjust various culture parameters to estimate their impact on cellular $O_2$ availability. It is accessible at https://www.lucidsci.com/notes?entry=oxygen_diffusion or as an open-source interactive notebook at https://observablehq.com/@lucid/oxygen-diffusion-and-flux-in-cell-culture.

#### Model of $O_2$ diffusion in medium of a culture well

As a cell monolayer consumes $O_2$ from culture media, more oxygen diffuses from air through the media. This movement of $O_2$ follows Fick's laws of diffusion.

We can describe the system with several key assumptions:

(1) Consumption of $O_2$ by the monolayer is uniform (i.e. no part of the well consumes more $O_2$ than another part)
(2) Diffusion only occurs vertically (i.e. the plastic walls of culture dishes are $O_2$-impermeant).
(3) Total OCR for the well is constant.
(4) The $O_2$ diffusion gradient has reached steady state (i.e. $[O_2]$ at any given vertical position is not changing over time).
(5) The media is static (i.e. no convection).

The solution to Fick's laws under these assumptions are:

$$C(z) = C_{sat} - \frac{J}{D} z$$

$$J = D \frac{C_{sat} - C(z)}{z}$$

- ***J*** is $O_2$ flux (which we term oxygen consumption rate, or OCR, in this study).
- ***z*** is depth below the top surface of the media.
- ***C***(z) is $O_2$ concentration as a function of depth.
- $\boldsymbol{C_{sat}}$ is the $O_2$ concentration at the air-media interface concentration (i.e. at ***z*** = 0).
- ***D*** is the $O_2$ diffusion coefficient for the media.

Both equations above are equivalent. The first form is useful in calculating $[O_2]$ as a function of distance from the air-media interface, given a certain OCR (i.e. - flux, J) rate. This equation emphasizes how $[O_2]$ decreases linearly with increasing distance from the air-media interface. The slope of decrease over distance is determined by the OCR (flux, J). The higher the OCR, the steeper the slope and therefore more significant the drop in [$O_2$]. The second form of the equation demonstrates how to calculate OCR is linearly proportional to the difference in [$O_2$] at the air-media interface and the [$O_2$] at a given depth.

With the above assumptions, all oxygen consumed by the cells occurs by diffusion vertically down through the media and therefore the flux (J) is equivalent to the OCR.

#### Calculating $O_2$ concentrations and determining maximum possible OCR

We can use the steady-state solution to Fick's equations to gain an understanding of how media volume impacts cellular [$O_2$]. Additionally, becausevFick's laws impose a limit to $O_2$ diffusion in a given period of time, we can also calculate the maximum OCR possible for various media depths. We do this by setting C(z) at the level of the cell monolayer to 0 (i.e. - all oxygen at cell monolayer is consumed).

To apply the steady-state solution, we need values for **D**, **$C_{sat}$**, and **z**. Based on reported values of $O_2$ diffusion in $H_2O$, we assume a diffusion coefficient **D** in media of 3e-3 $mm^2/s$ at 37°C [1]. Estimates of **D** vary in the literature, and are dependent on media composition, pH, temperature, and other factors. Thus, imprecision in the correct value of **D** for a given experimental situation may lend variability in the absolute [$O_2$] seen at the cell monolayer.

***z*** represents the position in the media relative to the top surface of the media. The maximum value is at the well bottom and is equal to the media height **H.** Assuming a perfectly cylindrical well, the media height is the length of a cylinder containing that amount of media volume so H = volume / ($\pi$ * $radius^2$). For typical 96-well plates, the radius is around 3.2 mm resulting in heights of 3.1 mm for 100 µL and 6.2 mm for 200 µL.

The dissolved $O_2$ concentration at the air-media boundary ***$C_{sat}$*** is determined by $O_2$ percent in the air, relative humidity, atmospheric pressure, temperature, and media $O_2$ solubility. For a cell culture incubator at 5% $CO_2$, high humidity, and ~760 mmHg, ***$C_{sat}$*** is ~175-204 µM [2].

To help with conversion between units of $O_2$ availability, at 37°C with high humidity, a rough conversion is that 1% $O_2$ in air equals 10 µM $O_2$ in media. This 1:10 conversion also holds when converting between dissolved $O_2$ in units of $O_2$% (v/v) and µM. The rough conversion rate is extrapolated from the following reference [2].

OCR is diffusion-limited when $O_2$ at the bottom of the well (at the cell monolayer) is zero, resulting in a steady-state max OCR of **$D * C_{sat} / H$,** where **H** is media height (i.e. the max OCR is inversely proportional to media volume).

#### Calculations with experiment data

The manuscript shows hypoxia at increased media volume. The Resipher $O_2$-sensing probes operate at 1-1.5 mm above the cell monolayer, and do not directly measure [$O_2$] at the cell monolayer. To approximate [$O_2$] the cells are experiencing, we apply the model above to OCR values reported for RPE cultures in the main manuscript. Using OCR values at 6 hours into the experiment as steady-state and assuming $\mathbf{C_{sat}}$ of 200 µM, estimated $O_2$ at cells varies across a range of media volumes, from around 110 µM (~11%) in 65 µL down to 25 µM (~2.5%) in 200 µL as shown in (**Supplementary Figure 3A**).

Additionally, we calculated the maximum possible OCR for each media volume and compared this with the reported OCR measurements. Unlike RPE cells cultured with 65 µL medium, RPE with 200 µL medium are operating at near maximum possible OCR (**Supplementary Figure 3B**).

The model we describe is employed in the linked calculator and interactive notebook. Using the calculator we can explore the relationship between media volume, OCR, and $O_2$ availability. In **Supplementary Figure 3C** we plot the relationship between media volumes and cellular $O_2$ availability at a constant OCR. In **Supplementary Figure 3D** we plot the relationship between OCR and cellular $O_2$ availability for a set of medium volumes. Additionally, different values for $\mathbf{C_{sat}}$ (e.g. for labs at high altitude or in $O_2$ controlled environments) can be used to calculate the effects on $O_2$ availability with various media volumes.

### Limitations

While this steady-state model is useful for understanding cell culture $O_2$ diffusion and calculating the impact of cell culture parameters on $O_2$ availability, there are a number of limitations to estimates of [$O_2$].

(1) $\mathbf{C_{sat}}$ is an important source of uncertainty. It is determined by many factors including ambient $O_2$ levels, atmospheric pressure, relative humidity, temperature, and media solubility. The effects of these factors are discussed extensively in [2]. Any bias in this value will impact the calculated $O_2$ at the cells. Under conditions where cells do not fully deplete culture medium of $O_2$, a 10 µM uncertainty in $\mathbf{C_{sat}}$ causes an identical 10 µM uncertainty in cellular $O_2$ availability.

(2) The $O_2$ diffusion constant **D**, also has a wide range of reported values. **D** is reliant on temperature, pH, media composition, and other factors. Reported values for **D** have varied as widely as 0.976–3.00 x$10^{-3}$ mm$^2$/s. [2]

(3) However, if we assume a true value of 2.5 - 3 x$10^{-3}$ mm$^2$/s, the uncertainties in **D** are equivalent to a +/- 10% uncertainty in media volume in determining cellular $O_2$ availability. [2]

(4) OCR can change rapidly, breaking steady-state assumptions. Dynamic, non-linear $O_2$ gradients that occur during these OCR changes cannot be modeled with the steady-state solution outlined above, and depending on how quickly changes in OCR occur, they may not be detected by the resipher instrument. Thus, estimates of cellular $O_2$ availability using our calculator should

be reserved for periods during cell culture when $O_2$ levels at the Resipher sensor are quite stable over time.

(5) The walls and bottom of the well plate are slightly oxygen permeable and cells or medium can get oxygen not only at the air-medium interface. This affects calculations of cellular $O_2$ availability. In addition, the walls of the cell culture plate are not cylindrical, but rather a conical frustum with the diameter tapering down from the top. The divergence from a cylindrical shape varies by manufacturer, and affects the reliability of the steady-state model. Additional extensive considerations that may affect calculations of cellular $O_2$ availability are detailed in [2].

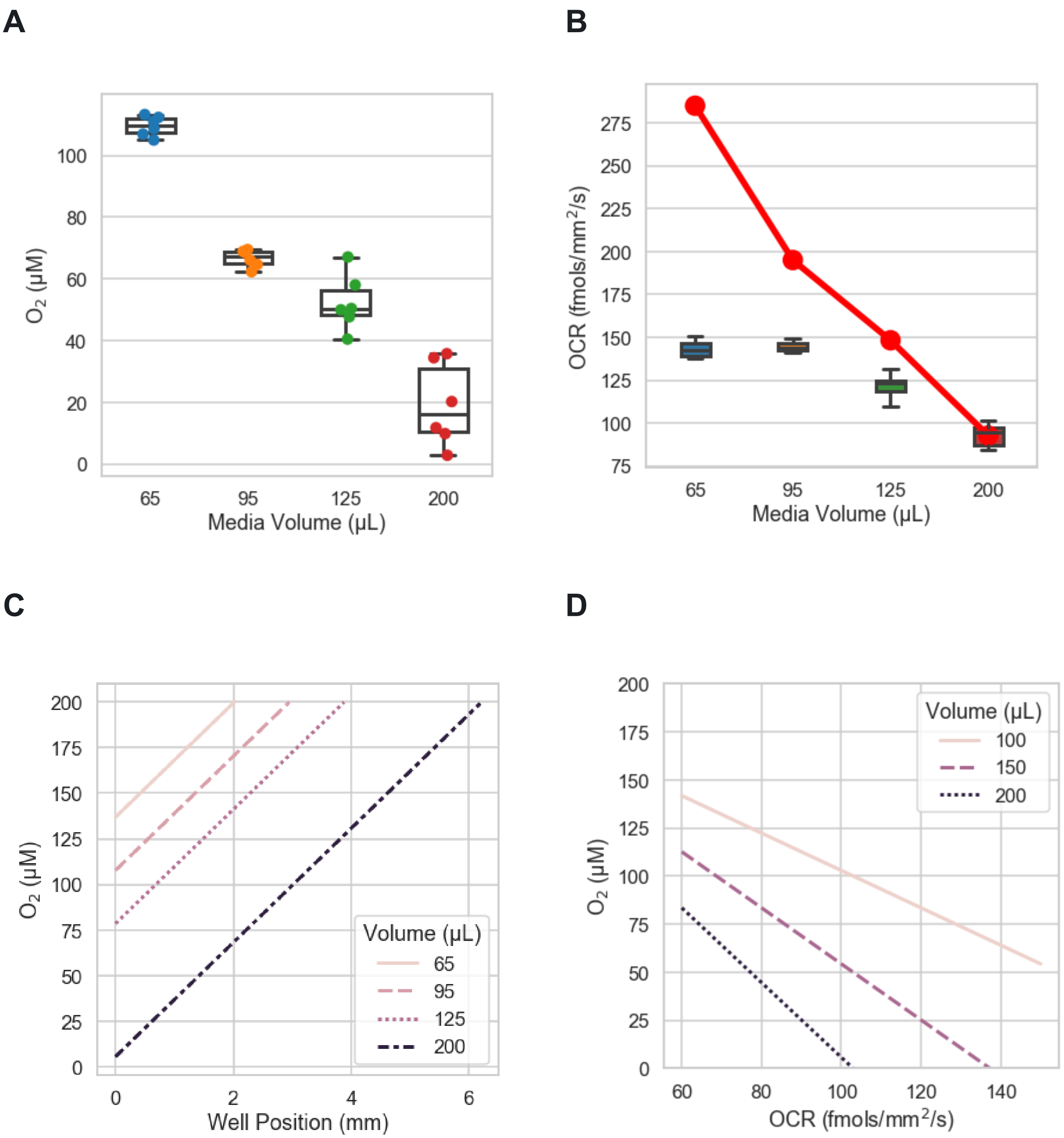


**Supplementary Figure 3. Calculations of *$O_2$ and OCR, and the effect of model parameters*** **(A)** Estimated $O_2$ available to cells as a function of medium volume, derived from experiment steady state OCRs after 6 hours. **(B)** OCRs (box and whiskers plots) at 6 hours compared with the maximum possible OCR for that medium volume (red). **(C)** Calculated $O_2$ levels as a function of medium depth for various media volumes. In each case, the hypothetical OCR is set at 100 fmols/mm$^2$/s. **(D)** Calculated $O_2$ availability at cells as a function of OCR for a set of fixed media volumes.
